# Neanderthals did not likely consume *Physcomitrium patens* – a model moss species

**DOI:** 10.1101/2022.01.26.472964

**Authors:** Fabian B. Haas, Raphael Eisenhofer, Mona Schreiber, Laura S. Weyrich, Stefan A. Rensing

**Affiliations:** University of Marburg, Plant Cell Biology, D-35043 Marburg, Germany; School of Biological Sciences, University of Adelaide, Australia; Department of Anthropology and Huck Institutes of the Life Sciences, Pennsylvania State University, University Park, USA

## Abstract

This manuscript is discussing the method of Weyrich *et al*., 2017, “Neanderthal behaviour, diet, and disease inferred from ancient DNA in dental calculus”. When studying the dietary profile of a Neanderthal specimen from El Sidrón cave (Spain) by sequencing ancient DNA present in calcified dental plaque (calculus) the authors identified a wide-range of potential food sources, including woolly rhinoceros, mushrooms, pine nuts, and moss – namely the less-than-abundant model species *Physcomitrium patens*. We doubted that Neanderthals were actually eating *P. patens*. By analyzing the ancient DNA reads using different mapping methods, we show likely a misinterpretation based on the previously used methods. The probability of Neanderthals eating *P. patens* is the same as eating rice or tomato. However, neither crop was grown in Europe at the time Neanderthals thrived.

## Did Neanderthals eat moss?

Deep sequencing of ancient DNA (aDNA) derived from digestive tracts and dental calculus has been used in recent years to infer dietary habits of past animals and humans. When studying the dietary profile of a Neanderthal specimen from El Sidrón cave (Spain) by sequencing ancient DNA present in calcified dental plaque (calculus), Weyrich *et al*. identified a wide-range of potential dietary food sources, including woolly rhinoceros, mushrooms, pine nuts, and moss ^1^. These species were identified using a MALTx approach, similar to BLASTx, which translates DNA sequences into proteins and runs comparisons to the NCBI non-redundant (nr) database (2014). Care was taken to ensure false mapping was not driving the results. For example, only DNA sequences > 50 bp were included in downstream analysis, and spurious mapping was filtered using the default last common ancestor (LCA) filter in MEGAN5 ^2^ prior to assessment. Additionally, assignments were ignored if the reference genome contained known human DNA contamination, the eukaryote was observed in a mock microbial metagenome, or the DNA was present in a laboratory blank control. Other contemporaneous aDNA analysis papers used similar approaches ^3^.

One particular dietary species of interest was a model moss species, *Physcomitrium patens* (previously known as *Physcomitrella patens*). While the range of this species makes consumption by Neanderthals a possibility, the ecology and lifestyle of this moss species contradict this finding. *P. patens* is an ephemeral annual that grows on disturbed, muddy ground ^4^. It has a small stature and occurs most commonly in late autumn, typically surviving for only a few weeks. Thus, the dietary use of this species seems unlikely. Yet, aDNA sequences in the Weyrich *et al*. paper mapped to the *P. patens* genome that was sequenced a decade ago ^5^ – a genome that has subsequently been refined and cleaned over the past ten years ^6^. Here, we aimed to further assess the identification of ancient DNA sequences that mapped to this genome.

## Do the number of hits found correlate with the sequence abundance in the database?

We applied several methods to re-assess whether there are *P. patens* reads in the El Sidrón 1 dental calculus DNA, using the above-mentioned stringent 50 bp cutoff. First, a Tera-BLASTn mapping against the NCBI GenBank nt database was carried out. In total, 12.6 million reads (39.4 %) of the aDNA present in the El Sidron dental calculus sample matched to 30,000 different genera within the GenBank nt data base (Table S1). The mapping of reads from El Sidrón 1 to *P. patens* was initially confirmed in this analysis. Yet, how accurate is this finding? A wealth of the Tera-BLASTn and BLASTn hits were also found for crops like rice, tomato, and corn (Tab. S2, Fig. S1). Critically, these species were not present in Europe during the Pleistocene and may have contaminated reference genome sequences, as discussed by Weyrich *et al*., highlighting potential problems with this approach. LCA filtration in MEGAN5 works to reduce false positive hits, as used in Weyrich *et al*., but the standards and settings for LCA analysis for this purpose this use remain poorly described.

Therefore, we selected eight species to further investigate using whole genome analysis: *Populus trichocarpa* (poplar, black cottonwood) and *Pinus koraiensis* (Korean pine) that were both mentioned by Weyrich *et al*., *Oryza sativa* Japonica (rice), *Zea mays* (corn), *Populus euphratica* (desert poplar), *Solanum lycopersicum* (tomato), *Solanum tuberosum* (potato), and *Nicotiana tabacum* (tobacco) that were chosen because they are popular models (hence well represented in the database) and have similar or larger genome sizes than moss. If we consider hits per Mbp genome size and per number of GenBank entries (Table S2), *P. patens* is on the lower side. While analysing the Tera-BLASTn results for the eight chosen species, we saw thousands of single read hits mapping exclusively to one specific species. For example, Tomato has 24,795 exclusive hits, whereas *P. patens* has 1,983 (Table S2). We then conducted short read mapping using GSNAP ^7^ against the *Physcomitrium patens* V3 genome ^6^ and these comparator species. We also subsequently extracted regions with a high read coverage, as such regions are less likely to represent spurious mappings. GSNAP mapped 59,413 reads to *P. patens*, with 68 % aligning to the nuclear genome and the remainder to the organellar genomes and rDNA sequences (i.e., 45 sequences to rDNA; Table S3). High coverage regions (HCR) were defined by a stringent minimum length of 50 bp and 10 x read coverage; 45 such regions where found using GSNAP in the *P. patens* V3 Genome (Table S4). Reverse BLAST searches with these HCR were carried out against GenBank nt, where 42 HCR led to BLAST hits (Table S5). Most hits mapped to the chloroplast genome (Table S5), probably due to the fact that the organellar genome is present in many copies per cell and much more conserved than the nuclear genome; such conserved stretches can be found across species boundaries. However, there are 97 HCR BLASTn hits mapping to the *P. patens* nuclear, chloroplast, and mitochondria genome (Table S6), but do they represent actual moss sequences? We performed the same experiment with rice (determining HCR, reverse BLASTing against GenBank) and found 1,542 hits to rice. Hence, unique hits from specific sources appear not to be a hallmark of true presence of this species’ DNA but may represent poor mapping. These hits probably are not only a function of the relative over-representation of these species in the database, but may be due to the relatively short aDNA read length (average 64.8 bp of the raw reads and 88.2 bp of the HCR).

More resent databases can help to increase the precision of such analyses ^8^. We reassessed sequences mapping to *P. patens* in the Weyrich *et al*. 2017 paper by using BLASTn against a newer NCBI nt database (November 2018). In this updated version of the NCBI nt database, the best BLASTn hit for the 10 unique *P. patens* sequences identified in the Weyrich *et al*. paper now more closely map to microbial species (n=8), including three oral bacterial species, *Actinomyces* oral taxon 414, *Desulfobulbus* ORNL TM7x, *Lactobacillus paracollinonides*, and potential contaminants species, such as *Aceitobacter* JWB. Some non-microbial hits (n=2), including *Ceratosolen solmsi* (fig wasp) and *Paulinella micropora* (amoeba), were also identified but are again unlikely to be true; importantly, *P. patens* was not identified. This finding suggests that cross mapping between microbes and true *P. patens* sequences may have arisen from poor mapping. Further, tools, not only databases, have also improved over time. MEGAN6 Community Edition (CE) now contains a new “weighted” LCA algorithm. The weighted LCA algorithm is more specific than the LCA algorithm used in MEGAN5. While using MEGAN6 CE and applying the weighted LCA algorithm to the El Sidrón 1 dental calculus DNA BLASTn results, we now find only 211 taxonomy classes, in contrast to 11,476 taxonomy classes found when using an unweighted LCA algorithm in MEGAN5, and *P. patens* was also not identified.

We next explored both poor mapping quality (i.e., short sequences spuriously mapping) and contamination in reference genomes. To do this, we constructed two simulated ancient microbial metagenomic data sets using 30 complete genomes from bacteria largely identified within Weyrich *et al*. (with the addition of E. coli) ^8^ compared to the same MALT database (2014 nr) using parameters reported in the Weyrich *et al*. 2017 paper. The 50 bp (log normal distribution; ancient DNA like) mock microbial sequences mapped to three eukaryotic reference species, while 100 bp (strict length; modern DNA like) mock microbial sequences mapped to four different eukaryote references, including two reference genomes known to contain human DNA contamination: *Medicago truncatula* (barrel clover) and *Caenorhabditis remanei* (nematode), and two others that were not found to be contaminated: *Pinus koraiensis* (Korean pine) and *Ricinus communis* (castor bean) ^9,10^. More stringent LCA parameters (weighted) to filter out poorly mapped sequences (i.e., those with < 90 % sequence identity to reference) reduced the total number of hits per reference, but did not remove all of them. The remaining hits had high average sequence identity to the references (> 95 %). This suggests that poor reference genome assemblies (i.e., that are contaminated with microbial DNA or otherwise poorly assembled) and poorly curated databases may also contribute to false positive results. Of note, the *P. patens* V3 genome was recently put through decontamination scrutiny ^6^ and assembled on chromosome scale, which may explain why sequences mapping to *P. patens* were not detected using this approach.

Overall, cross-mapping of microbial DNA sequences to eukaryotic references and contaminated reference sequences limit our ability to clearly verify *P. patens* as a Neanderthal food source. Therefore, we posit that it is unlikely Neanderthals were consuming *P. patens*, and that cleaner databases, decontaminated genomes, and more efficient mapping strategies for metagenomics data sets will need to be developed to facilitate effective eukaryotic DNA mapping from ancient metagenomic specimens in the future. Until these advancements occur, ancient metagenomic analyses of dietary food sources will benefit from interdisciplinary approaches ^11^.

## Acknowledgements

We are grateful for access to the de.NBI infrastructure (AG Goesmann, Gießen).

