## Supplementary Materials for "Neanderthals did not likely consume *Physcomitrium patens* – a model moss species"

**Supplementary Materials**
**Neandertals did not likely consume *Physcomitrium patens***

DNA sample
The DNA sample of Neanderthal El Sidrón 1 was downloaded at https://dx.doi.org/10.4225/55/584775546a409, section 2_Processed, sample 2NoAdapt_ELSIDRON1L7_lTACTG_rCTCGA_R1R2_Collapsed.fastq (https://www.oagr.org.au/api/v1/dataset_file/3086/download/2NoAdapt_ELSIDRON1L7_lTACTG_rCTCGA_R1R2_Collapsed.fastq). The processed sample, filtered by read length >= 50 bp, has 32,025,748 reads with a length distribution of 50 bp to 181 bp.

Full dataset Tera-BLASTn
The downloaded FASTQ file of Neanderthal El Sidrón 1 sample was split into 20 files of equal size. These 20 files were mapped against the NCBI GenBank nt database (version 2016-06-03) using Tera-BLASTn 9.0.0 on DeCypher 9.0.0.25 (http://www.timelogic.com/catalog/757/tera-blast). The E-value threshold was set to 0.1 and the maximal alignment value was set to 100. The Tera-BLASTn results of the 20 subfiles were merged into one file.

**Table S1: Results of the Tera-BLASTn mapping against the NCBI GenBank nt database.**
39.4 % of all DNA reads from the Neanderthal El Sidrón 1 sample yielded a hit in the GenBank nt database. These 12.6 million reads generated 150 million hits with around 30,000 genera involved.

| | >= 50 bp reads |
|---|---|
| Number of reads mapped to the NCBI GenBank nt database | 12,615,596 (39.4 %) |
| Number of reads with no hit | 19,410,152 (60.6 %) |
| Total number of Tera-BLASTn hits | 150,101,630 |
| Involved GenBank entries | 2.67 million |

**Table S2: Tera-BLASTn results of eight selected species to compare with *P. patens*.**
"Number of GenBank entries" represents the number of NCBI GenBank nt (version 2016-06-03) sequence entries for the nine species. "Tera-BLASTn hits" are all Tera-BLASTn hits found for the species in question. The number contains exclusively mapped reads as well as reads mapped to multiple organisms. The column "Exclusive hits" shows the number of hits mapped only to the species in the respective row. The last two columns show Tera-BLASTn nt hits divided by the genome size in Mbp and the number of GenBank entries, respectively. For *Pinus koraiensis* no genome size data were found and hence the average value for the genus Pinus was used (http://data.kew.org/cvalues/). *P. patens* has 11,666 hits, resulting from 10,216 reads. 1,983 reads/hits are mapping exclusively to *P. patens*. The average alignment length of these 1,983 hits is 25.4 bp (min. 23 bp, max. 46 bp), the average identity of the hits is 98.5 % (min. 89.1 %, max. 100 %) and the average query coverage is 40.5 % (min. 16.1 %, max. 88 %). 888 of the 11,666 hits match organellar/rDNA sequences (chloroplast: 717; mitochondrial: 133; ribosomal: 38).

| Common name | Species | Genome size [Mbp] | Number of GenBank entries | Tera-BLASTn hits | Exclusive hits | Tera-BLASTn hits / Genome size [Mbp] | Tera-BLASTn hits / Gb entries |
|---|---|---|---|---|---|---|---|
| Rice | *Oryza sativa* Japonica | 382 | 95,733 | 214,723 | 3,060 | 562 | 2.2 |
| Tomato | *Solanum lycopersicum* | 760 | 60,263 | 287,202 | 24,795 | 378 | 4.8 |
| Poplar | *Populus trichocarpa* | 417 | 54,157 | 47,146 | 4,364 | 113 | 0.9 |
| Poplar | *Populus euphratica* | 496 | 58,302 | 28,933 | 1,292 | 58 | 0.5 |
| Corn | *Zea mays* | 2,104 | 167,471 | 87,598 | 4,291 | 42 | 0.5 |
| Potato | *Solanum tuberosum* | 706 | 46,304 | 27,475 | 2,100 | 39 | 0.6 |
| Moss | *Physcomitrella patens* | 478 | 36,747 | 11,666 | 1,983 | 24 | 0.3 |
| Tobacco | *Nicotiana tabacum* | 3,643 | 107,070 | 3,647 | 59 | 1 | 0.0 |
| Pine | *Pinus koraiensis* | 26,430 | 2,458 | 124 | 1 | 0.005 | 0.1 |

A)

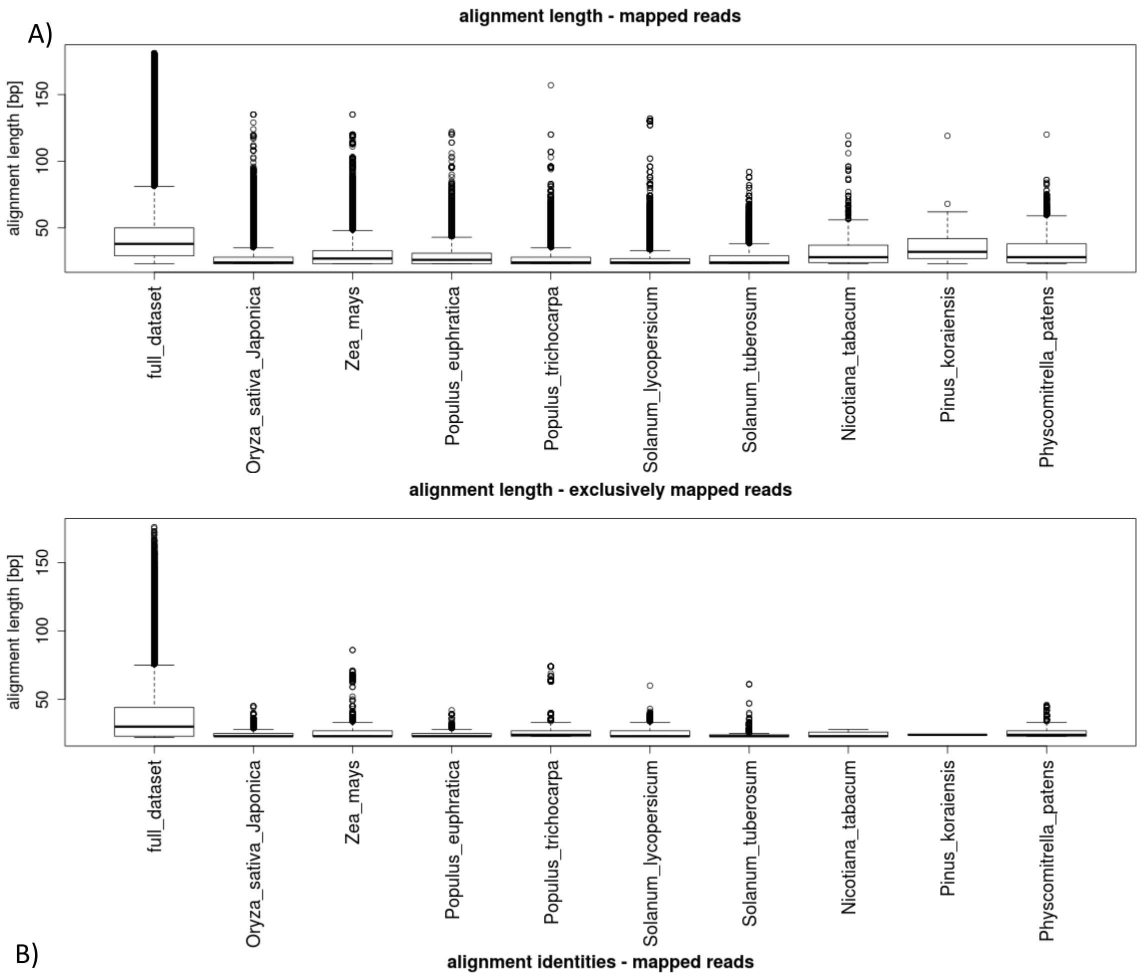
alignment length - mapped reads
alignment length [bp]
50
100
150
full_dataset
Oryza_sativa_Japonica
Zea_mays
Populus_euphratica
Populus_trichocarpa
Solanum_lycopersicum
Solanum_tuberosum
Nicotiana_tabacum
Pinus_koraiensis
Physcomitrella_patens
alignment length - exclusively mapped reads
alignment length [bp]
50
100
150
full_dataset
Oryza_sativa_Japonica
Zea_mays
Populus_euphratica
Populus_trichocarpa
Solanum_lycopersicum
Solanum_tuberosum
Nicotiana_tabacum
Pinus_koraiensis
Physcomitrella_patens

B)

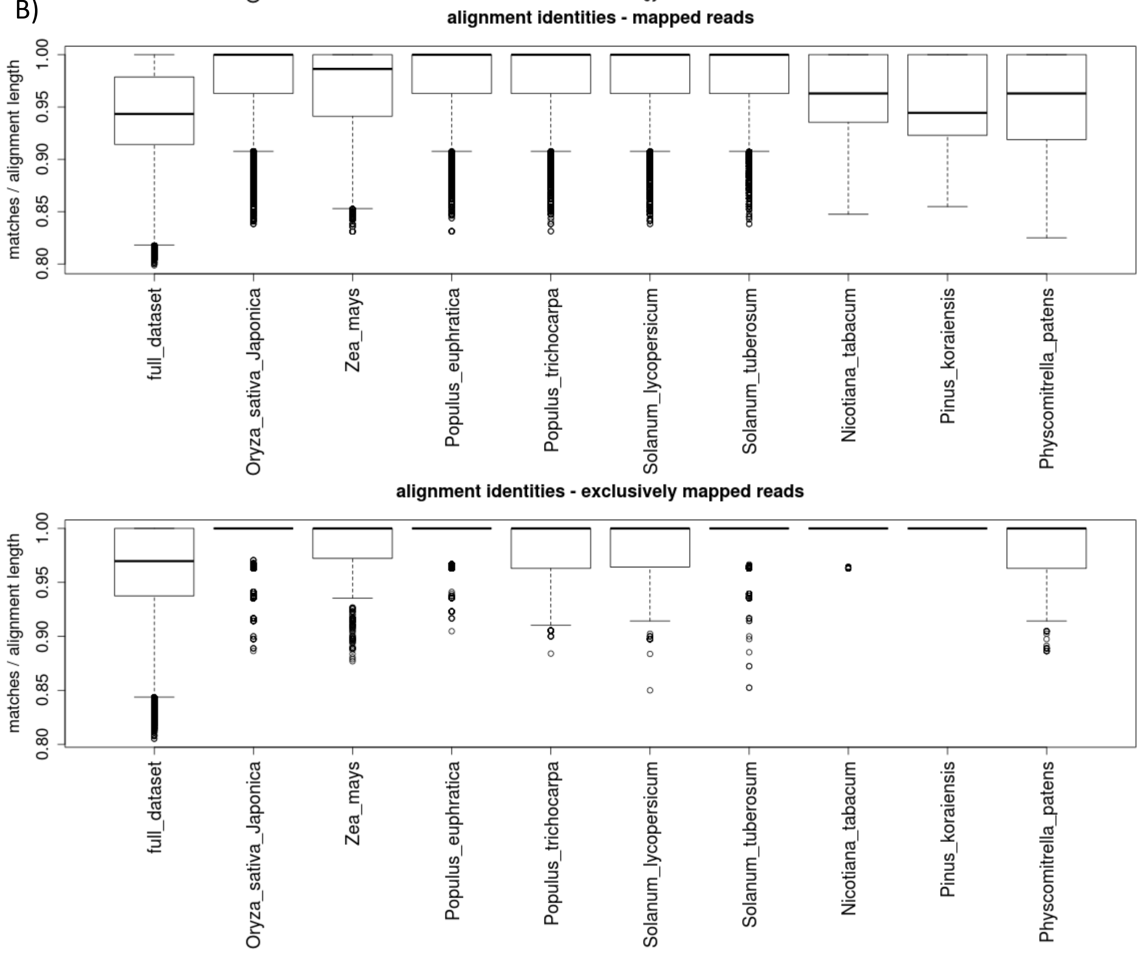
alignment identities - mapped reads
matches / alignment length
0.80
0.85
0.90
0.95
1.00
full_dataset
Oryza_sativa_Japonica
Zea_mays
Populus_euphratica
Populus_trichocarpa
Solanum_lycopersicum
Solanum_tuberosum
Nicotiana_tabacum
Pinus_koraiensis
Physcomitrella_patens
alignment identities - exclusively mapped reads
matches / alignment length
0.80
0.85
0.90
0.95
1.00
full_dataset
Oryza_sativa_Japonica
Zea_mays
Populus_euphratica
Populus_trichocarpa
Solanum_lycopersicum
Solanum_tuberosum
Nicotiana_tabacum
Pinus_koraiensis
Physcomitrella_patens

C)

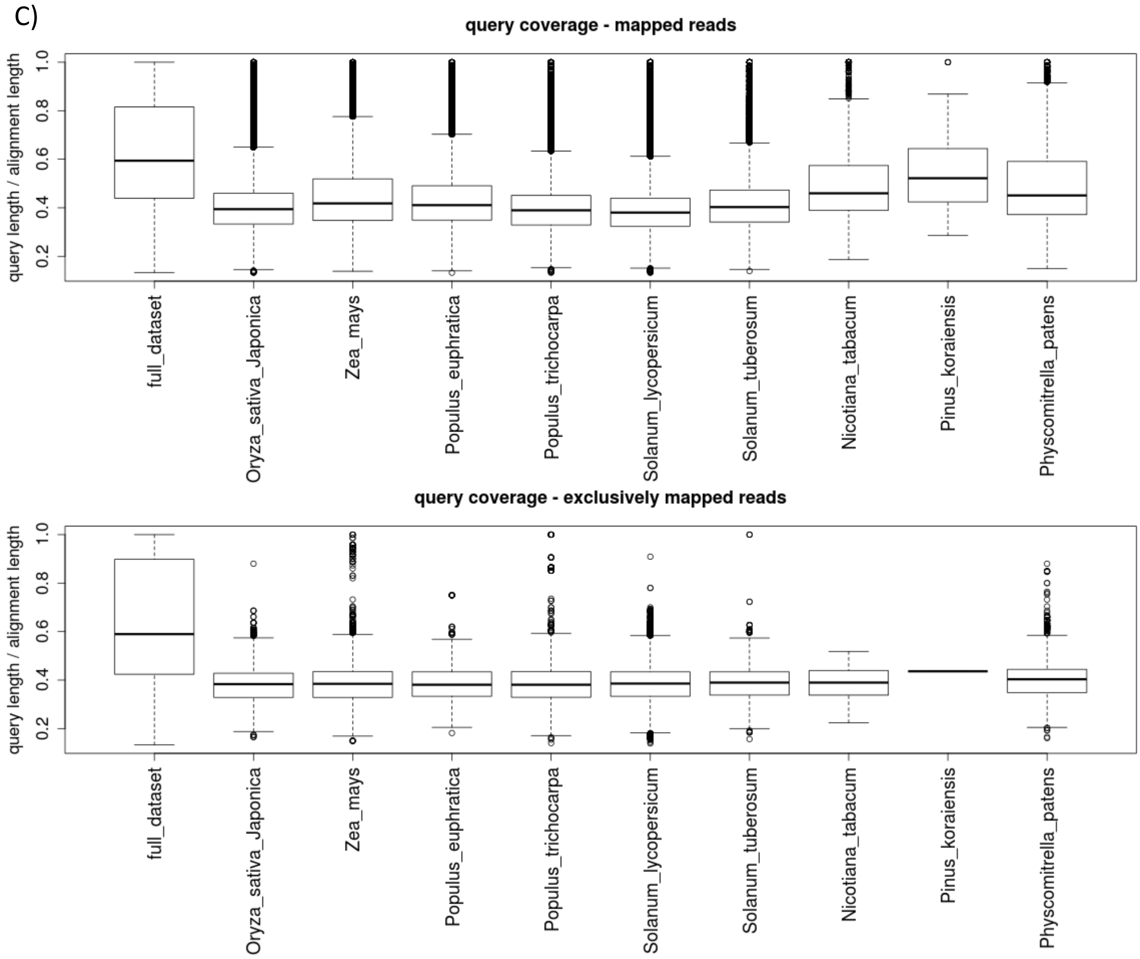


**Figure S1: Boxplots of the full dataset mapping result by Tera-BLASTn.**

Shown are alignment length (A), alignment identity (B) and query coverage (C).

The x-axis shows the results of the full dataset in the leftmost boxplot, followed by subsets of the nine species mentioned in Table S2. A) The y-axis represents the alignment length in bp for each hit. The maximum read length is <= 181 bp. B) On the y-axis the alignment identity (positive matches / alignment length) of the hits is plotted. The lowest value is 80%. C) The query coverage (query length / alignment length) is shown on the y-axis. All boxplots are shown for all Tera-BLASTn hits and the subset of the exclusively mapped hits. *P. patens* shows no obviously different distribution than the other species.

Full dataset short read mapping

The downloaded FASTQ file was mapped with the short read mapper GSNAP version 2016-11-07[1] against the *Physcomitrella patens* chloroplast (NCBI NC_037465) and mitochondrial (NCBI KY126309) genomes, the rDNA (HM751653, X80986, X98013) sequences and the *P. patens* nuclear genome V3[2]. All settings were kept in default mode. With the default GSNAP mismatch equation, 2 to 14 mismatches, depending on the read length, were allowed for this dataset.

**Table S3: Mapping result of the full Neanderthal El Sidrón 1 dataset mapped with GSNAP against the *P. patens* V3 nuclear genome, the genomes of chloroplast and mitochondria and the rDNA sequences.** 37,948 reads are mapping uniquely to the *P. patens* V3 nuclear genome. In total, 59,413 reads are mapping to the four genomic datasets. The multiple mapped reads can be found in the *P. patens* nuclear and chloroplast genomes. The mapping rate of 0.19 % is very low. 7,775 mappings are within *P. patens* v3.3 gene models.

| | Uniquely mapped reads | Multiple mapped reads |
|---|---|---|
| Chloroplast | 907 | 14,213 |
| Mitochondrion | 3,932 | 0 |
| rDNA | 45 | 0 |
| Nuclear genome | 37,948 | 2,368 |
| Sum | 42,832 | 16,581 |

High coverage regions (HCR)

Samtools depth version 1.3.1[3] was used to calculate the coverage of reads per nucleotide position. Using awk, highly covered read clusters were extracted. Threshold for a high coverage region (HCR) was a minimum length of 50 bp and a minimum coverage of 10 x per nucleotide. The sequences of the high coverage regions, found by GSNAP/Samtools, were extracted from the *P. patens* V3 reference genome as FASTA files and mapped with BLASTn (version 2.2.29+) against the NCBI GenBank nt database (version 2017-09-01). The E-value was set to 0.1 and the maximum alignment value was set to 5,000.

**Table S4: Number of high coverage regions with a minimum length of 50 bp and a coverage of 10 x.** Most of the 45 regions were found in the nuclear genome. The maximum coverage is 963 x and the highest average coverage is 252, both detected on the chloroplast genome. The length distribution is between 50 bp and 378 bp (in total 4,056 bp).

| | Number of regions | Max. region length [bp] | Average region length [bp] | Max coverage [x] | Average coverage [x] |
|---|---|---|---|---|---|
| Chloroplast | 23 | 378 | 101 | 963 | 252 |
| Mitochondrion | 16 | 129 | 90 | 411 | 127 |
| rDNA | 0 | 0 | 0 | 0 | 0 |
| Nuclear Genome | 6 | 63 | 56 | 37 | 21 |

**Table S5: Number of high coverage regions possessing BLASTn hits, and the number of BLASTn hits produced by these regions.** 42 of the 45 high coverage regions have hits against the NCBI GenBank nt database. In total, 252,255 hits were found. 157,927 (63 %) of all BLASTn hits were resulting from the chloroplast high coverage regions.

| | Number of HCC | Number of BLASTN hits |
|---|---|---|
| Chloroplast | 23 | 157,927 |
| Mitochondrion | 16 | 94,094 |
| Ribosome | 0 | 0 |
| Core Genome | 3 | 234 |

**Table S6: Number of BLASTn hits for the high coverage regions against the NCBI GenBank nt database, eight chosen species to compare with *P. patens*.**

The number of BLASTn hits and the ratio of BLASTn hit to genome size (see Table 2) is shown in column three and four. Rice and tomato harbour most hits relative to their genome size. None of the HCRs have an exclusive hit in any shown species. All of the 42 high coverage regions, which are mapping to the NCBI GenBank nt database, are mapping to *P. patens* (with 97 hits).

| Common name | Species | Number BLASTn hits | BLASTn hits / Genome size [MB] |
|---|---|---|---|
| Rice | *Oryza sativa Japonica* | 577 | 1.51 |
| Tomato | *Solanum lycopersicum* | 478 | 0.63 |
| Moss | *Physcomitrella patens* | 97 | 0.20 |
| Potato | *Solanum tuberosum* | 197 | 0.28 |
| Corn | *Zea mays* | 462 | 0.22 |
| Poplar tree | *Populus trichocarpa* | 72 | 0.17 |
| Poplar tree | *Populus euphratica* | 43 | 0.09 |
| Tobacco | *Nicotiana tabacum* | 170 | 0.05 |
| Pine | *Pinus koraiensis* | 18 | 0.0007 |

Rice genome

To determine whether other species would also yield exclusive hits we were tackling rice, the genome of which is approximately the same size as the moss genome, and which is not known to have been cultivated in Europe during the Pleistocene. The rice genome, GenBank assembly accession GCA_000005425.2, was used according to the methods “Full dataset short read mapping” and “High coverage regions (HCR)”. Using GSNAP, 1,363,331 (4.3 %) of the Neanderthal reads were mapping to the rice genome. 30,863 high coverage regions could be detected of which 96 regions had a BLASTn hit against the NCBI GenBank nt database.
